# A dCas9 Proximity Reporter for Investigating DNA Linear and Rotational Dynamics

**DOI:** 10.64898/2026.07.27.741122

**Authors:** Saba Parvez, Morgan C. Marsh, Shawn C. Owen, Randall T. Peterson

**Author notes:** equal contributions (Saba Parvez, Morgan C. Marsh).

## Abstract

The dCas9 system has rapidly been developed into many tools to explore different aspects of the human genome; the high binding specificity, coupled with the inactive nuclease enzyme, allows for precise recruitment of molecules to specific sequences of DNA. We sought to exploit these capabilities to create a tool for assessing the real-time proximity of two DNA sequences within a biological system. By incorporating aptamers into gRNAs, dCas9 molecules can be used to recruit the β9 or β10 strands of split-NanoLuc® to specific DNA sequences and quantify the proximity of those sequences based on their ability to complex with the luciferase fragment (Δ11S) and produce luminescence. While many tools exist to detect a single DNA sequence, this system is uniquely capable of assessing how two DNA sequences interact with each other. As expected, we found that the interaction of two dCas9 molecules was affected by their linear distance from each other on dsDNA. Surprisingly, we found that their interaction was also strongly influenced by rotational orientation, even for sequences that are close together in linear space. This finding indicates that dCas9 rotational alignment is an important consideration for designing dCas9 systems that target multiple DNA sequences simultaneously. Beyond the findings presented herein, we believe this DNA proximity detection tool has the potential to be adapted for applications involving the proximity and orientation of two DNA sequences.

## Introduction

Advances in CRISPR-Cas systems have provided new tools to modify and elucidate gene function. Among these, a catalytically inactive form of Cas9, dead Cas9 (dCas9), has been adapted for gene regulation,^1, 2^ biosensing,^3–5^ epigenetic modifications^6, 7^ and genomic imaging.^8, 9^ The possibilities using dCas9 are vast, with new tools and techniques rapidly being developed. Many of these tools focus on binding to a single site on DNA for specific detection or modification.

The human genome is complex, relying on both 3D and linear DNA organization as well as epigenetic modifications for gene regulation;^10, 11^ however, little is known about how the topography of the genome influences the expression of genes. It is postulated that not only DNA modifications^12, 13^ but also 3D orientation plays a role in gene regulation.^14, 15^ Currently there are multiple tools using dCas9 to look at epigenetic modifications and to physically alter these modifications to change gene regulation.^6, 7, 16–18^ Fewer tools exist to investigate the structural element of gene regulation. To this end, we developed a tool to better understand the proximity of different DNA sequences and the factors that influence their interactions in biological contexts.

Multiple tools have been developed using the dCas9 system that successfully detect single genomic loci with high precision.^19–25^ However, low signal-to-noise ratio remains an important limitation. Assays using fluorescent proteins fused to dCas9 are susceptible to photobleaching over the time of exposure or are limited by fluorescence intensity.^26^ These systems also have an always “ON” nature and rely on increased localization at the target site for accurate detection, which can lead to either false positive due to high background signal or false negatives if not enough fluorophore localizes. In a separate approach, dCas9 has been fused directly to split- luciferase proteins to detect single nucleotide polymorphisms (SNPs) in a gene.^25^ The luciferase system is specific and requires two dCas9 to bind in order to make a functional luciferase enzyme creating luminescent signal. This approach solves both the always “ON” and low sensitivity that can be seen with fluorophores. It resolves the high background issue as luminescence is only seen when two dCas9 molecules are bound. Additionally, the signal is substrate limited; therefore, low accumulation of the reporter can be addressed with higher substrate concentrations. Modified approaches include tandem RNA aptamers appended to the guide RNA (gRNA) that recruit fusion proteins consisting of an aptamer binding protein and a reporter. This allows multiple reporter proteins to bind the same region thereby increasing the overall signal.^21, 22^ Other methods employ proximity ligation and amplification techniques to increase the overall signal in cells for loci determination.^19, 20^ While all these methods have shown to be effective in detecting a single locus with high specificity, few have expanded to monitor the relative location of two genomic loci. Our system focuses on detection of two sequences in proximity.

Using unmodified dCas9, we developed a proximity labeling tool to detect neighboring sequences of interest (**Fig 1**). The system consists of dCas9, a pair of gRNAs modified with MS2 and PP7 RNA aptamers, the coat proteins MCP and PCP fused to β10 and β9 peptides, respectively, of a tri-part NanoLuc®, and Δ11S, the third part of the tri-part luciferase system. dCas9 is catalytically inactive but is easily programmed by gRNAs to bind to target sequences of interest with single nucleotide specificity.^27, 28^ The aptamers, PP7 and MS2, are validated for high protein binding specificity to the coat proteins, PCP and MCP, respectively.^29, 30^ Δ11S is an engineered version of NanoLuc® that requires complexation with β9 and β10, the two missing β-strands, to form an active enzyme.^31–33^ The β-strands need to be in close proximity for complementation with Δ11S to produce luminescence. Importantly, the complexation of the Nanoluc reporter is reversible^33^ which reduces non-specific signal due to spontaneous reporter complexation (**Supplementary Fig 1**). In the reporter assay, dCas9 complexes with gRNAs and binds to specified genomic targets. The RNA aptamers on the gRNAs recruit the fusion proteins β10-MCP and β9-PCP. If the two genetic loci are in proximity, the Δ11S complexes with the two β-strands creating an active enzyme complex that converts the provided substrate into luminescence. This approach provides a simple and robust readout to determine proximity between two separate DNA sequences.

**Fig 1.**
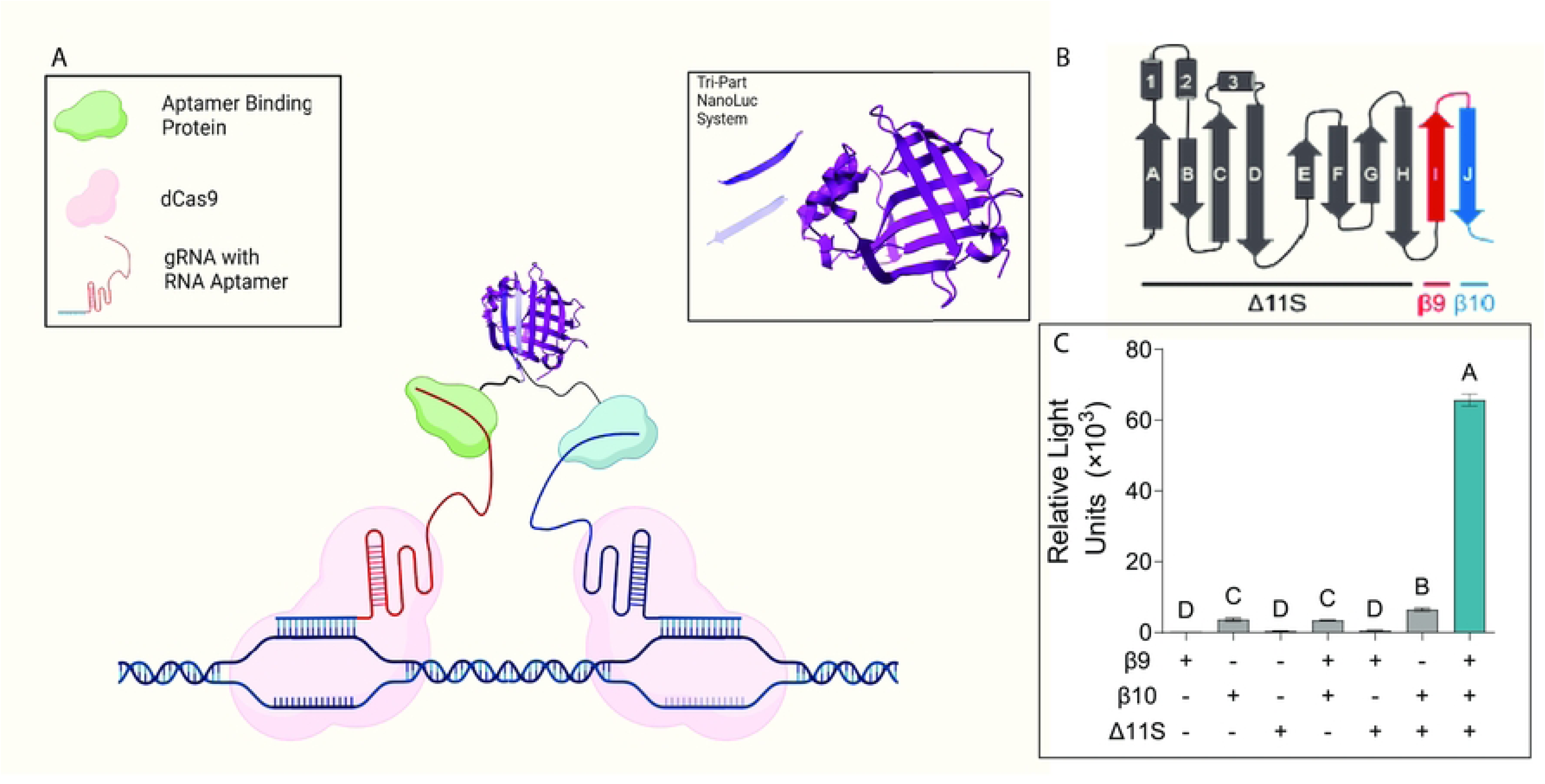
Components of the proximity reporter system. A: A proximity reporter system involving two dCas9 with two distinct gRNAs and a tri-part luciferase reporter. The gRNAs are extended on the 3’ end to include the RNA aptamers, MS2 and PP7. These aptamers recruit fusion proteins, β10-MCP and β9-PCP, containing the β9 and β10 strands of a trip-part NanoLuc® reporter, needed for recomplexation with the Δ11S portion. When the fusion proteins are in proximity, the β-strands can recomplex with the Δ11S making an active enzyme that converts substrate into a luminescent signal. **B:** Schematic indicating the different portions of the tri-part NanoLuc® in a linear representation. **C:** Simple graphic displaying luminescent activity is only seen when all three parts are added in vitro in excess. Data are presented as Mean ± SD (N =3). 1-way ANOVA with Brown-Forsythe and Welch was performed and compact lettering display used. Those that share a letter are not significant to each other (P>0.01).

## Methods

### Cloning for bacterial expression

All genes were cloned using ligase independent cloning in a pet21a vector for bacterial expression using primers indicated in **Supplementary Table 1**. The dCas9 was cloned out of Addgene plasmid # 72269 using dCas9-TEV-6xHis-fwd-1 and dCas9-TEV-6×His-rev-1 primers. An extension step was performed using dCas9-TEV-6xHis-fwd-2 and dCas9-TEV-6xHis-rev-2. The dCas9-TEV-6xHis-fwd-3 and dCas9-TEV-6×His-rev-3 primers were then used to add a TEV protease site and a 6x His tag to the C-terminus of dCas9. The resulting ‘megaprimer’ was introduced into empty pet21a plasmid linearized with EcoRI. Similarly, MCP and PCP were cloned out of Addgene plasmids #62330 and #66565, respectively using the indicated primers. Two extension steps were performed to add either the luciferase β9 or β10 fragment followed by a (GGGSG)3× linker on the N-terminus and a TEV protease site followed by a 6x His tag to the C-terminus of the aptamer binding proteins. All plasmid sequences were verified by Sanger sequencing (**Supplementary Table 1**).

### Cloning for mammalian expression

The MCP-β10 and PCP-β9 inserts were cloned into the pSICO vector (Addgene plasmids pJZC33 and pJCZC43 as templates) using ligation independent cloning with the primers and final insert sequence outline in **Supplementary Table 2**. Briefly, the fusion proteins were cloned in using the pSICO-lucb10, pSICO-lucb9 forward primers and the MCP-2×flag, PCP-2×flag reverse primers. Three extensions were made to generate the ‘megaprimers’, which were then introduced to the linearized pJZC33 and pJCZC43 vectors, respectively. The Δ11S was introduced into a pcDNA3 vector using primers outlined in Supplementary Table 2. The insert was made using the linker-Δ11S forward and Δ11S-pcDNA3 reverse primers and using pF1K-Δ11S as template^33^. Two subsequent extensions were made, and the insert was introduced to pcDNA3-dCas9-SV40-NLS (Addgene plasmid #100091) after linearizing with Kpn1 and Not1. The gRNA-2xMS2 and gRNA-2xPP7 sequences were amplified using pJZC33 or pJZC43 as the templates, respectively, and using bstXI-Alu-sgRNA forward and either sgRNA-XhoI or sgRNA-NotI reverse primers for introduction of specific restriction sites. The amplicon was digested with restriction enzymes and ligated in vectors containing the corresponding fusion proteins. Primers containing the specific gRNA sequences are outlined in **Supplementary Table 2**. All plasmid sequences were verified by Sanger sequencing.

### Protein production and purification

The pet21a plasmid containing the gene of interest was transformed into BL21-codon plus (DE3) cells. A single colony was inoculated in 5 mL Luria Broth (LB) media containing the selection antibiotic, kanamycin. The culture grew overnight at 37 °C and was added to 1 L of fresh LB-kanamycin media the next morning. The culture was grown to OD_600_ of 0.6-0.8 at 37 °C in a shaker at 200 rpm and induced with 250 µM IPTG for 3 h at 37 °C. Cells were harvested by centrifugation, flash frozen in liquid N_2_, and stored at -80 °C. For each expression, 3-4 g of cell pellet was obtained per liter of LB culture. The cell pellet was thawed on ice and resuspended with 5 mL lysis buffer/g of cell pellet. The lysis buffer contained 50 mM Na_2_HPO_4_ (pH 7.5), 150 mM NaCl, 5 mM Imidazole, 5 mM beta-mercaptoethanol (βME), 1 mM phenylmethanesulfonyl fluoride (PMSF). The cells were lysed by sonication and centrifuged at 20,000g for 30 minutes to remove any cell debris. The supernatant was incubated with 7 mL of HisPur cobalt resin for 1 h at 4 °C, after which the sample was loaded on a column and the flowthrough was discarded. The resin was washed under gravity with wash buffer (50 mM Na_2_HPO_4_, 300 mM NaCl, 10 mM Imidazole, 5 mM βME pH7.5). The bound protein was then eluted with elution buffer (50 mM Na_2_HPO_4_, 150 mM NaCl, 250 mM Imidazole, 5 mM βME pH 7.5. Elution fractions were collected every 2 mL. Fractions containing the protein, based off Nanodrop readings of the fractions, were combined and dialyzed overnight in dialysis buffer (50 mM HEPES, 100 mM KCl, 1 mM TCEP pH7.5) at 4 °C using a Slid-A-Lyzer^™^ G2 dialysis cassette 30 mL, 10 kDa molecular weight cut off (ThermoFisher Scientific® cat #87732). For dCas9, the dialyzed sample was then passed through an ion exchange column (5 mL SP FF) in Buffer A (50 mM HEPES (pH 7.5) and 100 mM KCl and eluted with a gradient of Buffer B (50 mM HEPES, 1 M KCl pH 7.5). Fractions containing the desired protein (determining by SDS-PAGE) were combined, the sample concentrated using AMICON 30 kDa MWCO concentrator and loaded on a size exclusion column (50 mM HEPES, 150 mM KCl, 10% glycerol, 1 mM TCEP) for final purification. Elution fractions corresponding to high 280/260 ratio were collected and concentrated to 10 mg/mL and stored in small aliquots in -80 °C. For the fusion proteins, a similar ion exchange and size exclusion methods were used except the buffers contained NaCl instead of KCl, and the samples after size exclusion were concentrated to 1 mg/mL in a 10 kDa MWCO concentrator. The protein sequences of the reporter components are listed in **Supplementary Table 3**.

### DNA templates for in vitro complementation assay

The DNA templates for in vitro complementation assay were generated in PAMin and PAMout orientation. To vary the distance between the proximal PAM sites, different number of nucleotides from an mcherry sequence (Addgene #62330) were inserted. For shorter sequences, a fill-in PCR using the indicated primers was used to generate the templates. For longer sequences, different lengths of the mcherry sequence were amplified using the indicated primers and using Addgene #62330 as the template plasmid. PCR products were cleaned up using Genejet PCR cleanup kit per manufacturer’s instruction and concentration was determined using Nanodrop. DNA sequences used in this study are listed in **Supplementary Table 4** and **Supplementary Table 5**.

### gRNA synthesis

DNA templates to generate single guide RNAs containing the RNA aptamer sequences 2xMS2 and 2xPP7 were generated by amplifying the aptamers from Addgene plasmids #62330 and #66565, respectively. An SP6 RNA polymerase site and a spacer sequence were added for in vitro transcription and target binding respectively (**Supplementary Table 1**). The 8xMS2, 8xPP7, 8xSirius_MS2, and 8xSirius_PP7 constructs containing an SP6 polymerase site and the spacer sequences were ordered as gene fragments from IDT. The templates were amplified using the indicated primers and the amplicon was column purified using the GeneJet PCR cleanup kit (ThermoFisher) following manufacture protocol. The sequences of DNA templates are listed in **Supplementary Table 6**.

In vitro transcription (IVT) was performed in RNAse free condition using a MEGAscript™ SP6 Transcription kit (ThermoFisher Scientific®, cat #AM1330) according to manufacturer’s guidelines. For each reaction of 20 μL, 6 pmol of DNA as well as 0.25 μL of RNAse inhibitor (ThermoFisher Scientific®; cat #EO0382) was used. The IVT sample was incubated at 37 °C overnight (∼16 h), after which the sample was treated with 1 μL Turbo DNAse for 15 min at 37 °C. Subsequently, the samples were cleaned up using an RNA Clean and Concentrator-5 (Zymo Research, cat #R1013) and eluted in 12 μL nuclease free water. The RNA concentration was determined using a Nanodrop (ThermoFisher Scientific®), RNA integrity assessed using gel electrophoresis, and the samples were then stored at −80 °C.

### Fluorescently labeling of gRNA for EMSA

The gRNA was diluted to 1.5 µM in nuclease free water then heated to 70 °C for 5 minutes. The gRNA was immediately placed on ice to cool. Using standard RNA ligation protocol, RNA was labeled with pCp-Cy5 (Jena Bioscience, NU-1706-CY5) appended to RNA with either the MCP or PCP aptamer sequence. Briefly, reactions were run with 1 µg gRNA, 1mM pCp-Cy5 RNA with 10 mM ATP and 10% DMSO in solution. T4-RNA ligase was added with the commercial buffer and reactions run for 2 hours at 16 °C.

### EMSA of protein DNA/RNA binding

In the binding buffer (20 mM HEPES, 250 mM KCl, 2 mM MgCl_2_ pH 7.5), dCas9 was diluted 1:1 with gRNA for a final concentration of 10 µM. The mixture was incubated at RT for 10 minutes. The dsDNA was added to the mixture for a final concentration of 23.5 nM and incubated at 37 °C for 10 minutes. Samples were loaded on a 0.8% agarose gel and run at 30V for 5-6 hours. The gel was incubated with SYBR safe DNA gel stain in 1× TAE buffer for 10 min, washed 3× with water and imaged. To assess gel shift upon coat protein binding to the aptamer, Cy5-labeled gRNAs were incubated at 70 ℃ for 5 minutes and rapidly cooled on ice to ensure proper aptamer folding. Samples contained 20 nM RNA and increasing ratios of the relevant fusion protein (β9-PCP for the PP7 aptamer and β10-MCP for the MS2 aptamer) in binding buffer (10 mM Tris (pH 7.6), 100 mM NaCl, 0.1 mM EDTA, 0.01 mg/ml tRNA, 50 μg/ml Heparin, 0.01% Nonidet P-40). These samples were then run on a 1.2% agarose gel (8.6 mM sodium tetraborate, 45 mM boric acid, pH 8.3).

### In vitro complementation assay

For the following proteins, 10× stocks were made. The β9-PCP (5 µM), β10-MCP (5 µM), and delta 11S (0.5 µM) were made in 1× PBSKT (137 mM NaCl, 10 mM Na_2_HPO_4_, 150 mM KCl, 0.1% Tween, pH 7.4) buffer. Ribo nuclear protein (RNP) complexes were made at 10× stocks of dCas9 bound to the different sgRNA (1 µM). RNPs were prepared by adding equimolar concentration of dCas9 and sgRNAs in 1× reaction buffer (20 mM HEPES, pH 7.5, 100 mM KCl, 5% glycerol, 1 mM DTT, 0.5 mM EDTA, and 2 mM MgCl_2_). The RNP mix was incubated at room temperature for 5-10 min. A 10× stocks of the DNA sequences (500 nM) were made in nuclease free water. The in vitro complementation reaction was set up in a white 96-well plate in a total volume of 50 µL. To set up, 10 µL of 5× PBSKT, 5 µL each of the 10x stocks of luciferase components, RNP and the DNA sequences were added to the wells. The samples were incubated at room temperature while gently shaking for 1 h. A 10× stock of coelenterazine (100 µM) was prepared in 1× PBST (from a stock of 2 mM in ethanol). Once sample had incubated for 1 h, 5 µL of the 10× coelenterazine was added to each well. Luciferase activity was measured every 5 min for 30 min at room temperature in a plate reader (Perkin Elmer VICTOR™ X3).

### In vitro complementation in 3D assay

All components were added simultaneously to the 96-well white plate at the following concentrations. In 1× PBSKT, the RNP was complexed at a 1:1 ratio of gRNA to dCas9 for a final concentration of 0.5 µM. Different concentrations of DNA were added with 25 nM final concentration of streptavidin for each reaction. For each fusion protein, 5 µM final concentration was bound and the whole complex was incubated for 2 hours with shaking. To the solution 0.5 µM of Δ11S with 10 µM of coelenterazine was added to the bound complex and luminescence measured.

### Identification of mammalian sites for complementation assay

The gRNAs were manually designed against consensus mammalian Alu sequence obtained from SINE Base^34^. Biostring package in R was used to identify the number of sites for each guide and the distance between two proximal gRNAs.

### Confocal microscopy

Cells were seeded in a glass 8-chamber slide and adhered overnight. Cells were transfected for 36 hours with a total of 500 ng of plasmid in the following ratios dCas9: Δ11S: β9-PCP: β10-MCP (3:1:3:5). Cells were then fixed with 4 % paraformaldehyde (PFA for 20 minutes at 4°C). The cells were then blocked with 0.2% Triton X-100, 3 % BSA in PBS for 1 hour at 37°C. Cells were then stained with α-flag (dCas9, Δ11S and fusion proteins contain flag tags) followed by a secondary antibody. Cells were also stained with DAPI (nuclear stain). Imaging was performed on a Zeiss LSM 700 laser scanning microscope and analyzed using Zeiss ZEN software.

### Luciferase complementation assay in cell culture

Cells were seeded at 5 x 10^4^ in a 96-well plate and incubated overnight at 37 °C with 5% CO_2_. Cells were then transfected using Mirus TransIT-2020 with different combinations of plasmids and control vectors for a total of 250 ng of plasmid DNA/well in the following ratios dCas9: Δ11S: β9-PCP: β10-MCP (3:1:3:5) with the addition of 2 ng Firefly luciferase vector. An empty pcDNA3 vector was used as a control in place of a protein vector of interest to keep the number of plasmids transfected the same. Cells were then incubated for 36 hours at 37 °C and 5 % CO_2_. Firefly and Nanoluc luciferase activity was measured using a Nano-Glo Dual-Luciferase Reporter Assay System (Promega, cat # N1630) according to manufacturer’s protocol. Briefly, Cells were lysed in passive lysis buffer and 25 µL of lysate was added to a white 96-well plate. To each well, 50 µL of Firefly substrate was added and luminescence measured. Next, 50 µL of NanoLuc® substrate was added to each well and luminescence was measured using a plate reader.

### Analysis of RNA structure using NUPACK

We used the web version of NUPACK software for the analysis of RNA secondary structure.^35^ RNA sequences (**Supplementary Table 6**) without the SP6 sites consisting of just the gRNA sequences modified with the tandem RNA aptamers were used as Input. The Minimum Free Energy (MFE) structure at 37 °C was generated.

## Results and discussion

To first test the efficacy of the reporter system *in vitro*, we recombinantly expressed individual protein components and purified them using affinity chromatography (**Fig 2A**). gRNAs were generated using in vitro transcription. We used an electrophoretic mobility shift assay (EMSA) to validate that the purified dCas9 bound to DNA in a gRNA-dependent manner (**Fig 2B**). Additionally, the fusion proteins β9-PCP and β10-MCP bound to gRNAs modified with two tandem repeats of the respective RNA aptamers (**Fig 2C**).

**Fig 2.**
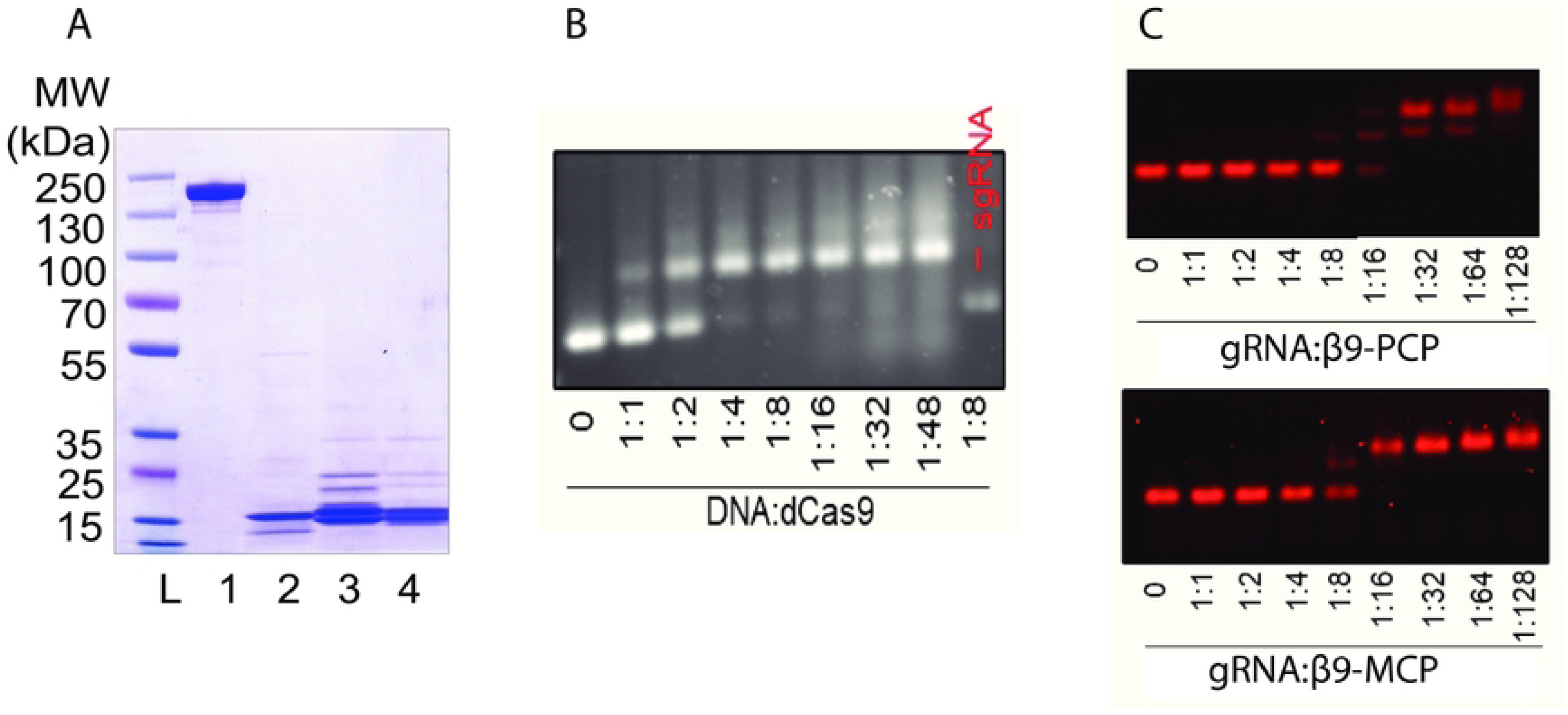
Production and validation of individual components of the proximity reporter. **A**: SDS-PAGE gel showing successful recombinant purification of dCas9 (lane 1; 158 kDa), Δ11S (lane 2, 17 kDa), β9-PCP (lane 3, 16.8 kDa), and β10-MCP (lane 4, 17.7 kDa). ‘L’ denotes molecular weight ladder. Expected molecular weights were calculated using the ExPasy ProtParam tool.^36^ **B**: An Electrophoretic Mobility Shift Assay (EMSA) showing dCas9-gRNA ribonucleoprotein complex successfully binds to DNA containing a dCas9 binding site. Titration of dCas9 shows a 1:4 ratio of DNA:dCas9 (23.5 nM of DNA) is sufficient to saturate DNA binding. No shift is seen without the gRNA highlighting the binding of dCas9 to the DNA is specific to the binding site. **C**: EMSA of fusion proteins binding to the gRNA aptamers. gRNAs contain two tandem repeats of the RNA aptamers, thus two higher molecular weight bands corresponding to the single and double occupied gRNA aptamers are seen with increasing the aptamer concentration of the fusion proteins.

Once the dCas9 and the fusion proteins were validated for successful binding, we then tested the reporter system *in vitro* to determine distance limitations and orientation of the dCas9 protein for optimal signal. We started by looking at dCas9 orientation in relation to the protospacer adjacent motif (PAM), a sequence in the DNA that is recognized by the dCas9 protein that sits downstream of the protospacer or targeting sequence. The PAM-IN orientation, where the two target sequences are on opposite DNA strands and the PAM sequences face one another was tested first (**Fig 3A**). A range of interspaced bases (4 base pairs (bp) – 711 bp) between the PAM sites were evaluated to determine the optimal distance for reporter complexation. The maximum signal was detected at 34 bp separation (11.56 nm) between the two proximal PAM sites and decreased when the two target sites were brought closer or moved farther apart from this optimal location. Interestingly, a strong periodicity was observed every 10 bp likely correlating with the degree of rotation on the dsDNA (**Fig 3A**). We repeated this study with the target sequences in PAM-OUT orientation where the two PAM sequences on the opposite DNA strands face away from each other (**Fig 3B**). The minimum number of intervening bases between the two PAMs in PAM-OUT orientation is 40 bp to ensure no overlap between the two protospacer sequences and to enable two dCas9s to bind the DNA simultaneously. Once again, a 5-10-fold signal increase was detected for most base intervals, indicating complexation. The highest signal was seen at 50 base pairs between the protospacers. Intriguingly, the same periodicity was observed suggesting that rotational orientation is important for optimal complexation.

**Fig 3:**
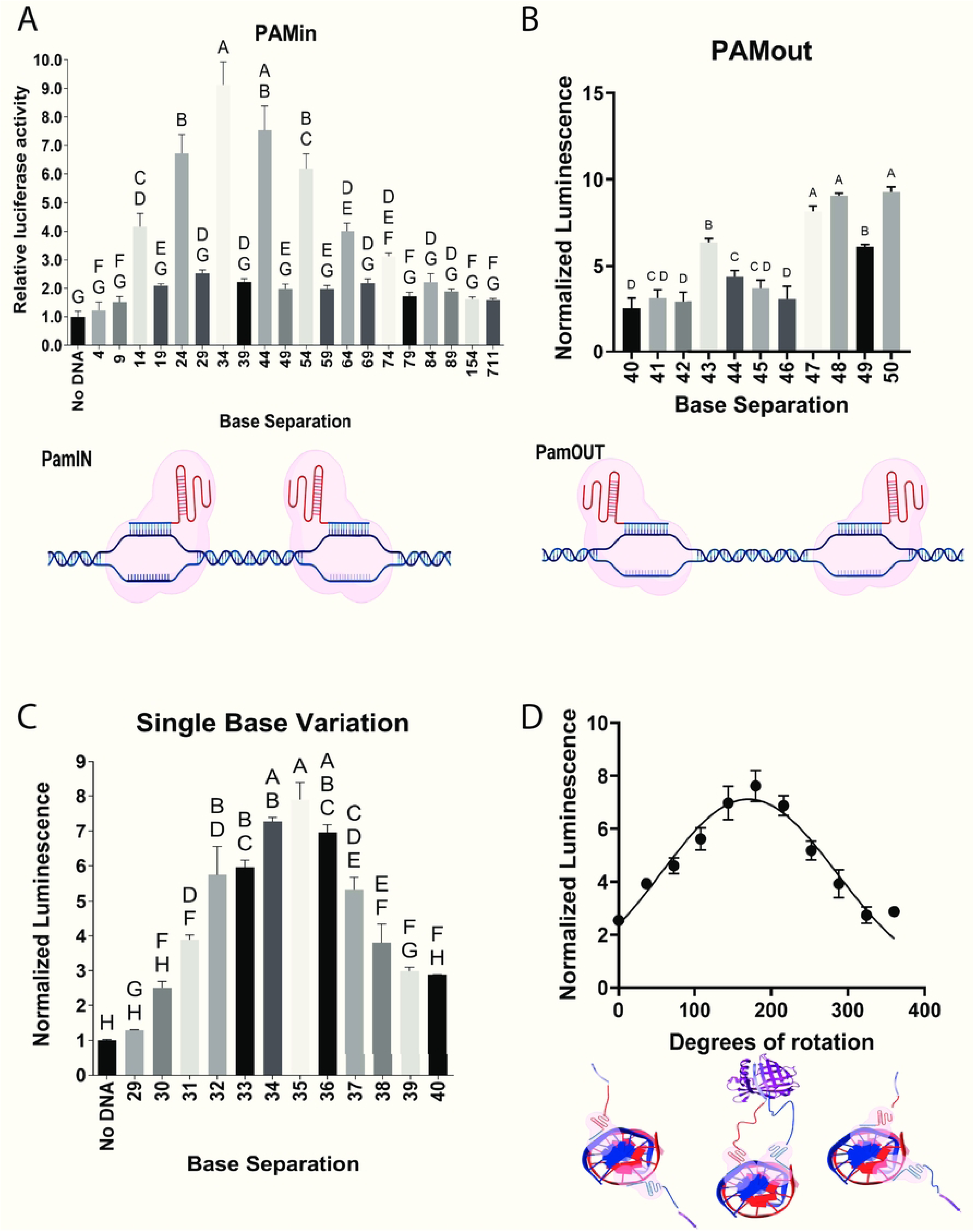
**Sensitivity of the proximity reporter to linear and rational distance of DNA *in vitro***. **A**: The effect of varying the distance between two target sites in PAM-IN orientation on the reporter activity. Distance is presented as the number of base pairs between the proximal PAM sites. The distance varied from 4 bp to 711 bp. A strong periodicity of ∼10 bp is observed in the PAM-in orientation. Sample with no DNA is used as control and all luciferase values are normalized against it. *Below*: Cartoon showing the target sites are in PAM-IN orientation. **B**: Relative luminescent signal at varying distance between the target sites in PAM-OUT orientation. Given that each target site is 20 bp in length, the minimum distance between two non-overlapping target sites in PAM-OUT orientation is 40 bp. *Below*: Cartoon showing the target sites are in PAM-OUT orientation C: Reporter activity with single base pair variation between two target sites in PAM-IN orientation **D**: Relative luminescent signal plotted against degrees of rotation of the DNA. When the two target sites are separated by 29 bp, the reporter constructs are on opposite sides of the DNA helix and are out of phase with each other. A single base pair addition changes the rotational orientation by around 36 degrees. At 34 bp gap (5 bp addition and a consequent 180° rotational change) between the two target sites, the reporter components are in phase resulting in more complexation and a higher luciferase signal. The reporter activity decreases with additional distance due to reporter components shifting to out of phase orientation *Below*: A 3D model of a DNA double helix shows that at a distance of 29 bp between the target sites in PAM-IN orientation, the fusion proteins recruited by the dCas9 and gRNAs are out of phase from each other. A 5 bp addition brings the fusion proteins in phase resulting in increased complexation. For statistical analysis, a 1-way ANOVA with Brown-Forsythe and Welch was run with compact lettering display indicating no significance (P>0.01) if a letter is shared between samples. A gaussian fit was added to the degrees of rotation. Data is presented as Mean ± SEM (N=3).

We then took a closer look at the periodicity of the system by measuring luminescence at single base pair intervals (**Fig 3C**) in the PAM-IN orientation and were able to determine the highest signal was seen when the dCas9 pair is in phase (**Fig 3D**). The design of the system includes multiple flexible linkers to facilitate complexation, which is seen at all degrees of rotation; however, the greatest increase in signal correlates with a rotation orientation that brings the pair into alignment. The periodicity and signal intensity relationship likely corresponds with the turn of a DNA double helix, as the gRNA will bind and conform to the double helix shape. The rotational specificity was an unexpected finding, indicating rotational orientation is an important factor to consider in determining proximity.

We tried various strategies to boost the signal of the proximity reporter. We varied the linker length between the gRNA and the RNA aptamer. Addition of the linker sequences did not perturb RNA folding or their ability to bind the fusion proteins (**Supplementary Fig 2**). Moderate increase in signal was seen with a longer linker length between the gRNAs and the RNA aptamers (**Fig 4A**). The greatest increase in signal was seen with a 5-base linker between the gRNAs and the RNA aptamers. Increasing the linker length by 20 bases gave the same signal as no increase indicating there are optimal distances for complexation. While we were able to see a moderate increase in signal by adjusting linker length, rotational orientation still contributed the most to the overall signal. Additionally, we increased the number of tandem RNA aptamers to eight to determine if higher signal would be seen with more reporter recruitment and consequent complexation. Intriguingly, when testing reporter activity with gRNAs modified with eight tandem repeats of the RNA aptamer in the tetraloop, no signal was seen (**Supplementary Fig 3**). While the addition of aptamers in the tetraloop does increase the overall number of fusion proteins bound (**Supplementary Fig 3**), it likely inhibits the complexation of the luciferase components. Although previously validated in cells,^37^ it is possible that tandem repeats of the aptamers in the tetraloop disrupts proper gRNA folding and/or impairs binding to dCas9 *in vitro*. We then moved the aptamer sites to the end of the gRNA, rather than in the tetraloop region, to determine if the placement of the aptamer was causing the lack of complexation. An EMSA displayed complete binding at a ratio of 1:64 sgRNA to fusion protein; multiple bands can be seen indicating multiple fusion proteins are binding (**Fig 4B**). Aptamer placement at the end of the gRNAs resulted in luciferase activity, however, the signal intensity with eight tandem aptamers still yielded an 8-10-fold signal change, similar to the original construct that only had two tandem aptamers (**Fig 4C**). Further optimization by titrating different components of the reporter did not result in any measurable increase in signal intensity (**Fig 4C**). Thus, it is likely that activity is limited by how many fusion proteins can bind simultaneously and still align to make active complexes. The system shows detection of two genes’ proximity, evidenced by the small changes to signal under multiple optimizations conditions.

**Fig 4:**
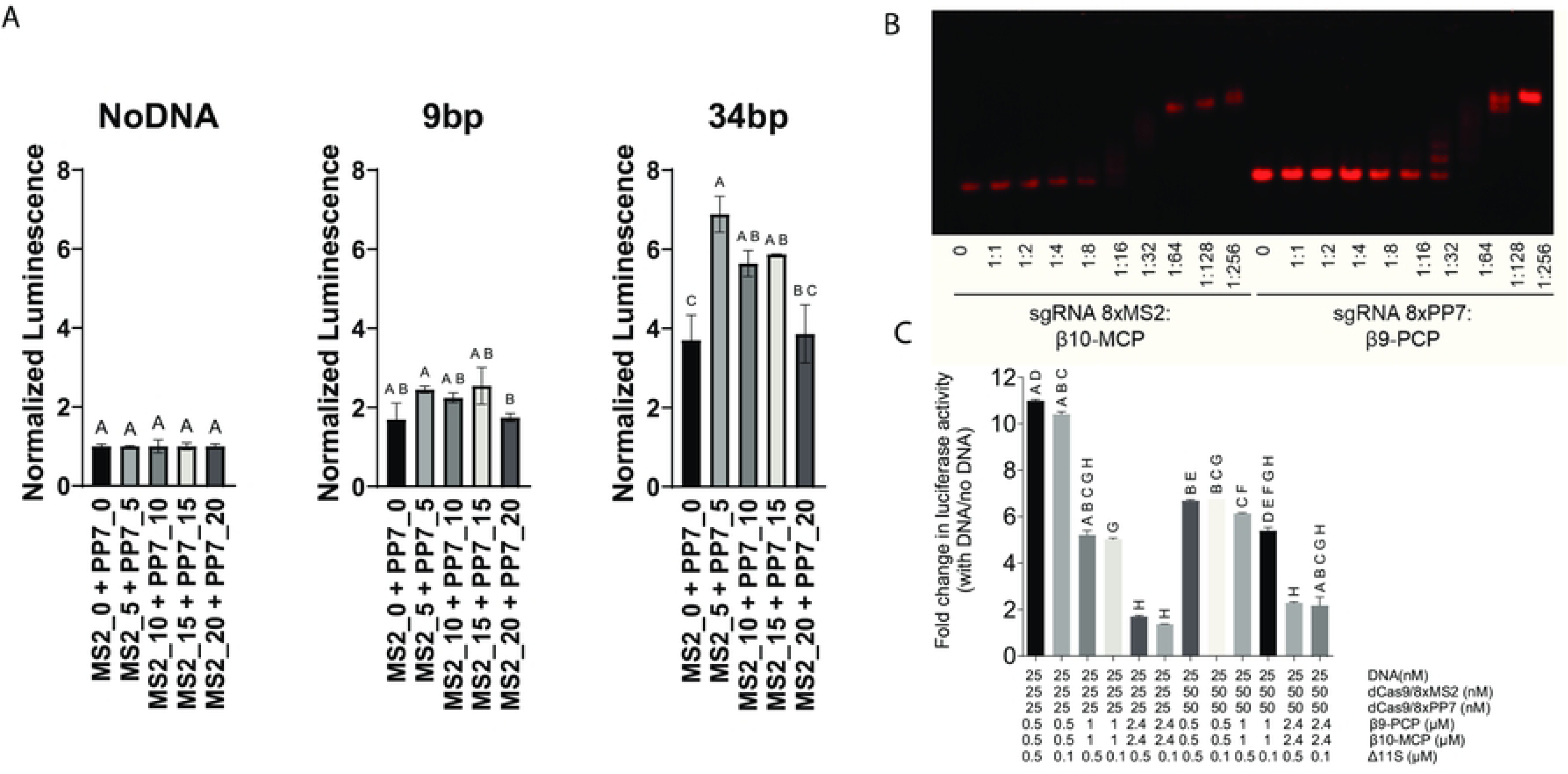
**Optimization of the proximity reporter**. **A**: Luminescent signal increases with longer linker length between the gRNA and the RNA aptamer. The numbers indicate the length of the linker. MS2_0 and PP7_0 are the standard gRNA-aptamer sequences with no linker between the gRNA and the aptamer. The greatest increase in signal is seen with a 5-base linker length but decreases with increasing linker length. The orientation of the DNA still seems to be the greatest contributing factor to proximity detection. **B**: An EMSA of fusion proteins binding with gRNAs modified with eight tandem RNA aptamer at the end of the gRNA sequence. Titration of the fusion proteins results in increasing number of tandem RNA aptamers being occupied as seen with multiple higher molecular weight bands. **C**: Luciferase activity with the eight tandem repeats of the RNA aptamer displays a 10-fold increase in signal over no DNA control. Additional optimization was done by varying the concentrations of each individual component. Increasing the concentrations of the fusion proteins and the aptamer sequences increases random complexation and background signal, resulting in a decreased fold change in luciferase activity. For statistical analysis, 1-way ANOVA with Brown-Forsythe and Welch was performed and compact lettering display used. Those that share a letter are not significant to each other (P>0.01). Data is presented as Mean ± SEM (N = 3) for all samples.

After validating that the reporter can successfully detect two proximal loci on a single piece of linear DNA, we moved to testing if the reporter could detect induced proximity when two loci on different DNA strands are brought together. This experiment aimed to mimic proximity between two loci in a 3D space. Thus, we biotinylated two separate stands of DNA containing the dCas9 binding sites (**Fig 5**). We added the biotin modification to either the 5’ or the 3’ ends of the DNA. All the individual components of the reporter system were added to a mix. Finally, streptavidin was added to induce proximity between the DNA strands and reporter activity was measured. Albeit small, a measurable increase in luciferase signal was seen when the 5’-biotin modified DNA was brought in proximity upon streptavidin addition, suggesting the proximity reporter may also detect proximal DNA sequences on separate DNA strands (although non-specific, streptavidin-mediated concentration of the components cannot be excluded). The increase in signal observed is smaller than in the linear format, indicating some optimization may be needed for improved sensitivity in 3D. While the exact distance between the two dCas9 proteins are difficult to determine in the 3D space, the detection of luminescent signal indicated our system still works in a non-linear system.

**Fig 5:**
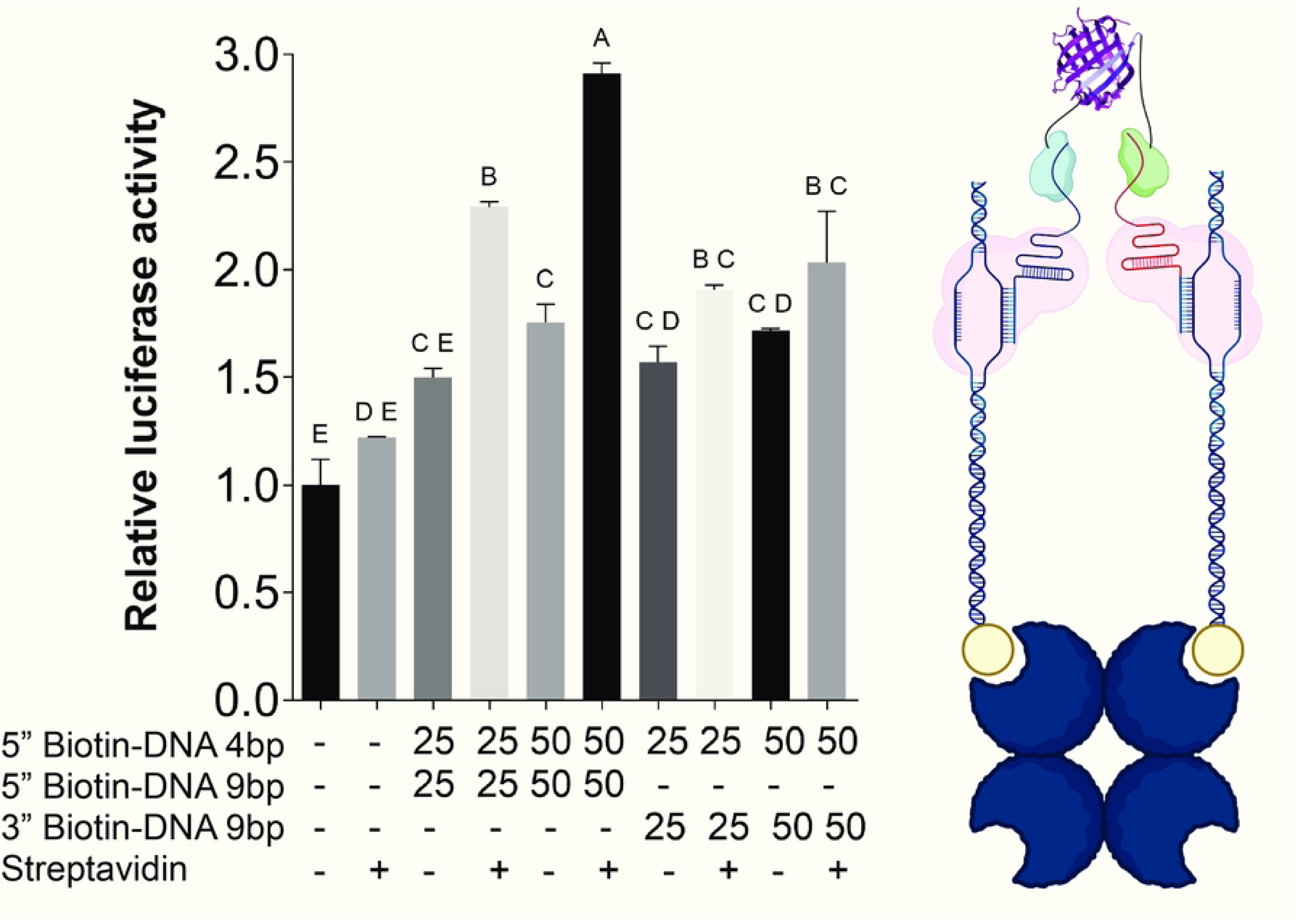
**Reporter activity with induced proximity in a 3D setting**. A 1.5-fold increase in luciferase signal is observed when two loci on separate pieces of DNA (DNA 1 and DNA 2) are brought together by induced proximity. The loci were on DNA that was biotinylated either on the 5’ or 3’ ends. Streptavidin was added to the assay mix to induce proximity between the DNA strands. Cartoon of a streptavidin tetramer. Only two bound DNA molecules are shown. Statistics were performed using a 1-way ANOVA with Brown-Forsythe and Welch. The compact lettering display indicates any bar that shares a letter is not significant (P>0.01). Data is presented as Mean ± SEM (N = 3)

Finally, we tested the reporter in mammalian cells. We targeted the reporter to Alu elements, transposable short DNA sequences that are in abundance, and measured the reporter’s sensitivity. Telomeres are useful for proof-of-concept detection of reporter systems because they are in abundance and at set locations on the chromosome. We decided to focus on Alu elements to determine if detection could occur within the chromosome but still ensuring signal with the high abundance. Alu elements are interspersed with genes and access is also subject to epigenetic modifications, making Alu elements a better test system to determine if detection of single gene proximity could be accomplished. We selected several pairs of target sites containing the NGG PAM (required for dCas9 binding) within Alu elements that are separated by 34 bp, the optimal distance for reporter complexation identified through in vitro assay. A computational search was used to determine the frequency of the target sites within the human genome and the distances between the gRNA pairs in PAM-IN orientation (**Fig 6A**). Based on this search, we selected 3 pairs of gRNAs that are present in differing frequencies within the human genome. Although Alu elements are conserved, variation within the sequences is also observed (**Fig 6A**). Nonetheless, the selected gRNA pairs were abundant in the human genome and were separated by the optimal distance in PAM-IN orientation indicating we should see good reporter activity in cells. We transfected HEK 293T cells with plasmids expressing each of the reporter components. We first performed confocal microscopy to check expression and localization of each protein component in the proximity detector. Imaging indicated good expression of all the components and sufficient localization to the nucleus (**Fig 6B and *inset***). Next, we co-transfected cells with plasmids expressing the protein components. The plasmid expressing the fusion proteins also expressed gRNAs modified with the RNA aptamer under a murine U6 promoter. Excitingly, we observed an 8-10-fold increase in luminescent signal when all the components were co-transfected (**Fig 6C).** Additionally, we observed that the signal intensity varied linearly with the number of paired target sites in the human genome. Importantly, very little background signal was seen when any individual component of the reporter was missing. For example, transfection of a control vector which lacks gRNAs expression resulted in the complete loss of luminescence demonstrating that the expression and complexation of the entire reporter is needed for the observed signal. Thus, these data illustrate the potential for detecting proximity between DNA sequences in cells, opening the possibility of several in cell applications. The proof-of-concept in cells focuses on known repeat elements, but the high signal suggests it may be possible to detect proximity between sequences that are less abundant, perhaps even as few as tens of copies per cell with optimization, although the exact limit of detection remains to be determined experimentally.

**Fig 6:**
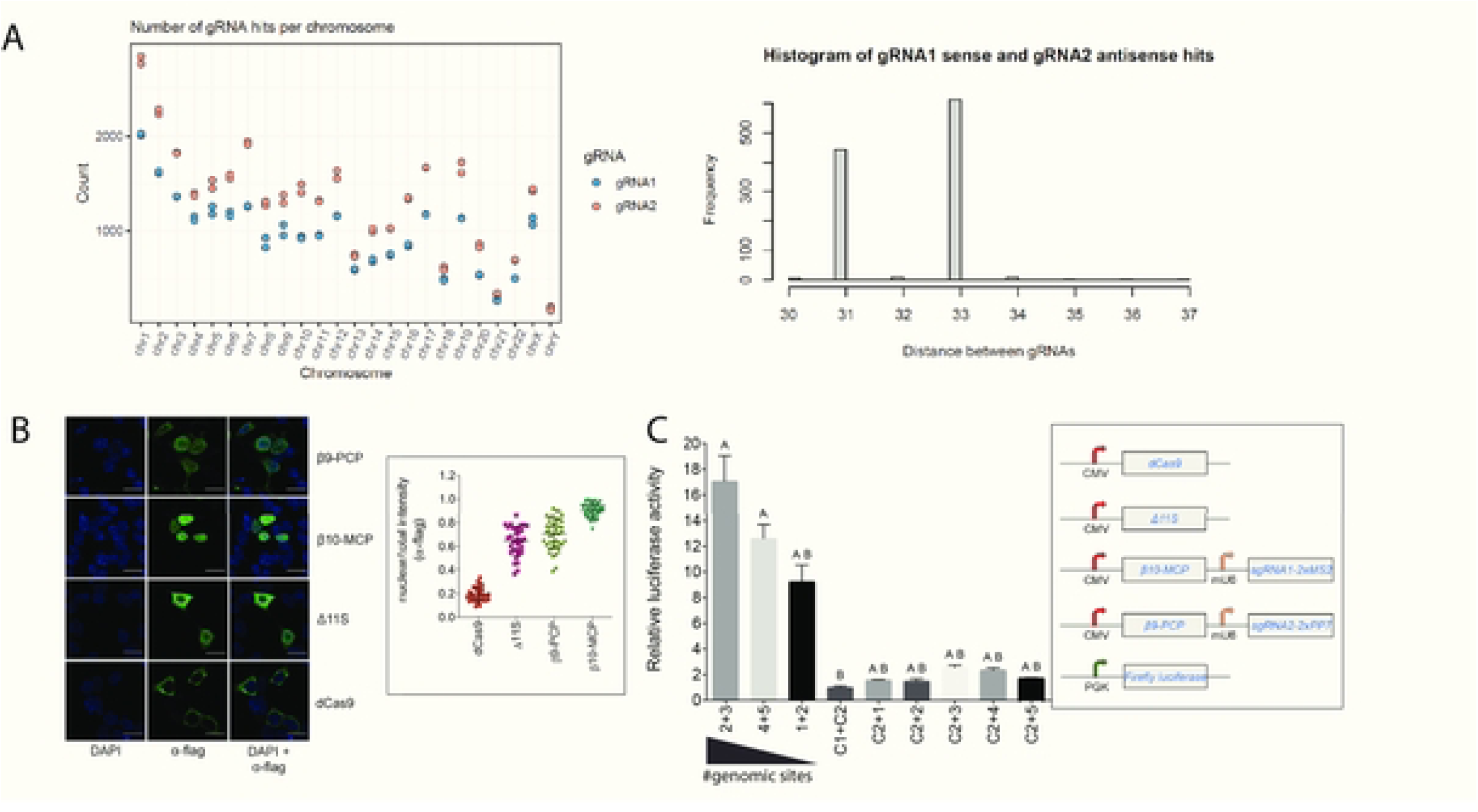
Proximity reporter testing in cells A: Frequency of a pair of gRNAs (gRNA1 and gRNA2) on the mammalian chromosome. *Inset*: Majority of gRNA1 and gRNA2 pairs are separated by 31 or 33 bp in PAM-IN orientation. **B**: Confocal imaging shows expressed protein in the nucleus (β9-PCP, β10-MCP) and in both the cytoplasm and nucleus (Δ11S, dCas9). The colocalization of constructs in the nucleus is sufficient to allow for site-directed complementation. *Inset*: Nuclear/total fluorescence quantitation in individual cells. **C**: Co-transfection of plasmids caused an increase in luminescent signal depending on how many genomic targets were present. gRNAs are labeled as 1-5. Control vectors that express the fusion proteins β10-MCP and β9-PCP without any gRNAs are labeled as C1 and C2, respectively. Co-expression of gRNA2 and gRNA3 pair gave the greatest signal because it has the highest frequency count in the genome. No signal was seen when one component was substituted with a control vector (C1 or C2) that expresses a non-targeting gRNA.

## Conclusions

We have developed a CRISPR-based proximity reporter by combining dCas9 with a tri-part split-luciferase system and demonstrated its ability to detect proximity between genomic loci *in vitro* and in cells. Using two dCas9-RNA complexes molecules to target different DNA sequences, we were able to determine the influence of linear distance, rotational distance, and PAM orientation on the interaction between two DNA loci. Surprisingly, we saw a strong rotational dependency for signal, a factor often ignored when performing CRISPR-based gene manipulation. This system displays the single nucleotide specificity that has been shown in other methods,^21–24^ with the addition of a second target detection and rotational information to give insight into 3D structure on linear DNA. Additionally, the ability to detect signal in 3D space indicates the potential of the system for determining gene proximity on histones or other 3D chromosomal elements. Given the current challenges in tracking real-time changes in chromosomal reorganization,^40^ specifically during different cell stages,^41^ this tool could potentially be adapated to investigate proximity of DNA sequences during dynamic cellular processes. In sum, these studies suggest that CRISPR-based proximity detectors like the one described here may be useful tools for multiple applications in the gene regulation and gene editing fields, while highlighting the critical importance of rotational orientation for dCas9 binding and interaction on DNA.

## Acknowledgements

We would like to acknowledge Chelsea Herdman and Joe Yost for determining the frequency of Alu elements for detection through this system.

## Supporting information

**S1 Table. Primer Sequences for Bacterial Expression plasmid cloning**

**S2 Table. Primer Sequences for Mammalian Expression plasmid cloning**

**S3 Table. Protein Sequences**

**S4 Table. DNA Sequences for the PAM-IN Orientation**

**S5 Table. DNA Sequences for the PAM-OUT Orientation**

**S6 Table. DNA Sequence for the gRNA Templates**

**S1 Fig. Comparison of the 2-component and 3-component split NanoLuc systems**

**S2 Fig. RNA aptamer modified gRNA structures and gels their binding with fusion proteins**

**S3 Fig. In vitro complementation assay with gRNAs modified with eight tandem RNA aptamers in the tetraloop**

**Supplementary Fig 4. Full gels images**

## Notes

### Competing Interest Statement

The authors have declared no competing interest.

## References

1. Lu, A.; Wang, J.; Sun, W.; Huang, W.; Cai, Z.; Zhao, G.; Wang, J., Reprogrammable CRISPR/dCas9-based recruitment of DNMT1 for site-specific DNA demethylation and gene regulation. Cell Discovery 2019, 5 (1), 22.

2. Savell, K. E.; Bach, S. V.; Zipperly, M. E.; Revanna, J. S.; Goska, N. A.; Tuscher, J. J.; Duke, C. G.; Sultan, F. A.; Burke, J. N.; Williams, D.; Ianov, L.; Day, J. J., A Neuron-Optimized CRISPR/dCas9 Activation System for Robust and Specific Gene Regulation. *eneuro* 2019, 6 (1), ENEURO.0495-18.2019.

3. Uygun, Z. O.; Yeniay, L.; Gi Rgi, N. S. F., CRISPR-dCas9 powered impedimetric biosensor for label-free detection of circulating tumor DNAs. Anal Chim Acta 2020, 1121, 35–41.

4. Jiang, H.; Zhu, X.; Jiao, J.; Yan, C.; Liu, K.; Chen, W.; Qin, P., CRISPR/dCas9-based hotspot self-assembling SERS biosensor integrated with a smartphone for simultaneous, ultrasensitive and robust detection of multiple pathogens. Biosensors and Bioelectronics 2025, 270, 116974.

5. Gao, X.; Tsang, J. C. H.; Gaba, F.; Wu, D.; Lu, L.; Liu, P., Comparison of TALE designer transcription factors and the CRISPR/dCas9 in regulation of gene expression by targeting enhancers. Nucleic Acids Research 2014, 42 (20), e155–e155.

6. Policarpi, C.; Munafò, M.; Tsagkris, S.; Carlini, V.; Hackett, J. A., Systematic epigenome editing captures the context-dependent instructive function of chromatin modifications. Nature Genetics 2024, 56 (6), 1168–1180.

7. Rajaram, N.; Kouroukli, A. G.; Bens, S.; Bashtrykov, P.; Jeltsch, A., Development of super-specific epigenome editing by targeted allele-specific DNA methylation. Epigenetics & Chromatin 2023, 16 (1), 41.

8. Clow, P. A.; Du, M.; Jillette, N.; Taghbalout, A.; Zhu, J. J.; Cheng, A. W., CRISPR-mediated multiplexed live cell imaging of nonrepetitive genomic loci with one guide RNA per locus. Nature Communications 2022, 13 (1), 1871.

9. Geng, Y.; Pertsinidis, A., Simple and versatile imaging of genomic loci in live mammalian cells and early pre-implantation embryos using CAS-LiveFISH. Sci Rep 2021, 11 (1), 12220.

10. Cremer, T.; Cremer, C., Chromosome territories, nuclear architecture and gene regulation in mammalian cells. Nature Reviews Genetics 2001, 2 (4), 292–301.

11. Puck, T. T.; Krystosek, A.; Chan, D. C., Genome regulation in mammalian cells. Somatic Cell and Molecular Genetics 1990, 16 (3), 257–265.

12. Cheung, P.; Allis, C. D.; Sassone-Corsi, P., Signaling to Chromatin through Histone Modifications. Cell 2000, 103 (2), 263–271.

13. Zhang, Y.; Reinberg, D., Transcription regulation by histone methylation: interplay between different covalent modifications of the core histone tails. Genes & Development 2001, 15 (18), 2343–2360.

14. Bonev, B.; Cavalli, G., Organization and function of the 3D genome. Nature Reviews Genetics 2016, 17 (11), 661–678.

15. Lanctôt, C.; Cheutin, T.; Cremer, M.; Cavalli, G.; Cremer, T., Dynamic genome architecture in the nuclear space: regulation of gene expression in three dimensions. Nature Reviews Genetics 2007, 8 (2), 104–115.

16. Brocken, D. J. W.; Tark-Dame, M.; Dame, R. T., dCas9: A Versatile Tool for Epigenome Editing. Curr Issues Mol Biol 2018, 26, 15–32.

17. Bhattacharjee, G.; Gohil, N.; Siruka, D.; Khambhati, K.; Maurya, R.; Ramakrishna, S.; Chu, D. T.; Singh, V., CRISPR-dCas9 system for epigenetic editing towards therapeutic applications. Prog Mol Biol Transl Sci 2023, 198, 15–24.

18. Gemberling, M. P.; Siklenka, K.; Rodriguez, E.; Tonn-Eisinger, K. R.; Barrera, A.; Liu, F.; Kantor, A.; Li, L.; Cigliola, V.; Hazlett, M. F.; Williams, C. A.; Bartelt, L. C.; Madigan, V. J.; Bodle, J. C.; Daniels, H.; Rouse, D. C.; Hilton, I. B.; Asokan, A.; Ciofani, M.; Poss, K. D.; Reddy, T. E.; West, A. E.; Gersbach, C. A., Transgenic mice for in vivo epigenome editing with CRISPR-based systems. Nature Methods 2021, 18 (8), 965–974.

19. Liang, Y.; Wu, S.; Han, W.; Wang, J.; Xu, C.; Shi, J.; Zhang, Z.; Gao, H.; Zhang, K.; Li, J., Visualizing Single-Nucleotide Variations in a Nuclear Genome Using Colocalization of Dual-Engineered CRISPR Probes. Analytical Chemistry 2022, 94 (34), 11745–11752.

20. Zhang, K.; Deng, R.; Teng, X.; Li, Y.; Sun, Y.; Ren, X.; Li, J., Direct Visualization of Single-Nucleotide Variation in mtDNA Using a CRISPR/Cas9-Mediated Proximity Ligation Assay. Journal of the American Chemical Society 2018, 140 (36), 11293–11301.

21. Wang, S.; Su, J.-H.; Zhang, F.; Zhuang, X., An RNA-aptamer-based two-color CRISPR labeling system. Scientific Reports 2016, 6 (1), 26857.

22. Shao, S.; Zhang, W.; Hu, H.; Xue, B.; Qin, J.; Sun, C.; Sun, Y.; Wei, W.; Sun, Y., Long-term dual-color tracking of genomic loci by modified sgRNAs of the CRISPR/Cas9 system. Nucleic Acids Res 2016, 44 (9), e86.

23. Zhang, Y.; Wang, Y.; Xu, L.; Lou, C.; Ouyang, Q.; Qian, L., Paired dCas9 design as a nucleic acid detection platform for pathogenic strains. Methods 2022, 203, 70–77.

24. Xu, X.; Luo, T.; Gao, J.; Lin, N.; Li, W.; Xia, X.; Wang, J., CRISPR-Assisted DNA Detection: A Novel dCas9-Based DNA Detection Technique. The CRISPR Journal 2020, 3 (6), 487–502.

25. Heath, N. G.; Segal, D. J., CRISPR-Based Split Luciferase as a Biosensor for Unique DNA Sequences In Situ. In Fluorescence In Situ Hybridization (FISH): Methods and Protocols, Haimovich, G., Ed. Springer US: New York, NY, 2024; pp 285–299.

26. Zhang, H.; Guo, P., Single molecule photobleaching (SMPB) technology for counting of RNA, DNA, protein and other molecules in nanoparticles and biological complexes by TIRF instrumentation. Methods 2014, 67 (2), 169–176.

27. Myers, S. A.; Wright, J.; Peckner, R.; Kalish, B. T.; Zhang, F.; Carr, S. A., Discovery of proteins associated with a predefined genomic locus via dCas9–APEX-mediated proximity labeling. Nature Methods 2018, 15 (6), 437–439.

28. Wang, C.; Qu, Y.; Cheng, J. K. W.; Hughes, N. W.; Zhang, Q.; Wang, M.; Cong, L., dCas9-based gene editing for cleavage-free genomic knock-in of long sequences. Nature Cell Biology 2022, 24 (2), 268–278.

29. Bertrand, E.; Chartrand, P.; Schaefer, M.; Shenoy, S. M.; Singer, R. H.; Long, R. M., Localization of ASH1 mRNA Particles in Living Yeast. Molecular Cell 1998, 2 (4), 437–445.

30. Chao, J. A.; Patskovsky, Y.; Almo, S. C.; Singer, R. H., Structural basis for the coevolution of a viral RNA–protein complex. Nature Structural & Molecular Biology 2008, 15 (1), 103–105.

31. Kim, S. J.; Yao, Z.; Marsh, M. C.; Eckert, D. M.; Kay, M. S.; Lyakisheva, A.; Pasic, M.; Bansal, A.; Birnboim, C.; Jha, P.; Galipeau, Y.; Langlois, M.-A.; Delgado, J. C.; Elgort, M. G.; Campbell, R. A.; Middleton, E. A.; Stagljar, I.; Owen, S. C., Homogeneous surrogate virus neutralization assay to rapidly assess neutralization activity of anti-SARS-CoV-2 antibodies. Nature Communications 2022, 13 (1).

32. Dixon, A. S.; Schwinn, M. K.; Hall, M. P.; Zimmerman, K.; Otto, P.; Lubben, T. H.; Butler, B. L.; Binkowski, B. F.; Machleidt, T.; Kirkland, T. A.; Wood, M. G.; Eggers, C. T.; Encell, L. P.; Wood, K. V., NanoLuc Complementation Reporter Optimized for Accurate Measurement of Protein Interactions in Cells. ACS Chemical Biology 2016, 11 (2), 400–408.

33. Dixon, A. S.; Kim, S. J.; Baumgartner, B. K.; Krippner, S.; Owen, S. C., A Tri-part Protein Complementation System Using Antibody-Small Peptide Fusions Enables Homogeneous Immunoassays. Scientific Reports 2017, 7 (1).

34. Vassetzky, N. S.; Kramerov, D. A., SINEBase: a database and tool for SINE analysis. Nucleic Acids Res 2013, 41 (Database issue), D83-9.

35. Zadeh, J. N.; Steenberg, C. D.; Bois, J. S.; Wolfe, B. R.; Pierce, M. B.; Khan, A. R.; Dirks, R. M.; Pierce, N. A., NUPACK: Analysis and design of nucleic acid systems. Journal of Computational Chemistry 2011, 32 (1), 170–173.

36. Gasteiger, E.; Hoogland, C.; Gattiker, A.; Duvaud, S. E.; Wilkins, M. R.; Appel, R. D.; Bairoch, A., Protein Identification and Analysis Tools on the ExPASy Server. In The Proteomics Protocols Handbook, Walker, J. M., Ed. Humana Press: Totowa, NJ, 2005; pp 571–607.

37. Ma, H.; Tu, L.-C.; Naseri, A.; Chung, Y.-C.; Grunwald, D.; Zhang, S.; Pederson, T., CRISPR-Sirius: RNA scaffolds for signal amplification in genome imaging. Nature Methods 2018, 15 (11), 928–931.

38. Hunt, J. M. T.; Samson, C. A.; Rand, A. D.; Sheppard, H. M., Unintended CRISPR-Cas9 editing outcomes: a review of the detection and prevalence of structural variants generated by gene-editing in human cells. Human Genetics 2023, 142 (6), 705–720.

39. Höijer, I.; Emmanouilidou, A.; Östlund, R.; van Schendel, R.; Bozorgpana, S.; Tijsterman, M.; Feuk, L.; Gyllensten, U.; den Hoed, M.; Ameur, A., CRISPR-Cas9 induces large structural variants at on-target and off-target sites in vivo that segregate across generations. Nature Communications 2022, 13 (1), 627.

40. Takata, H.; Masuda, Y.; Ohmido, N., CRISPR imaging reveals chromatin fluctuation at the centromere region related to cellular senescence. Scientific Reports 2023, 13 (1), 14609.

41. Nagano, T.; Lubling, Y.; Várnai, C.; Dudley, C.; Leung, W.; Baran, Y.; Mendelson Cohen, N.; Wingett, S.; Fraser, P.; Tanay, A., Cell-cycle dynamics of chromosomal organization at single-cell resolution. Nature 2017, 547 (7661), 61–67.

42. Mei, Q.; Huang, J.; Chen, W.; Tang, J.; Xu, C.; Yu, Q.; Cheng, Y.; Ma, L.; Yu, X.; Li, S., Regulation of DNA replication-coupled histone gene expression. Oncotarget 2017, 8 (55), 95005–95022.

43. Prado, F.; Jimeno-González, S.; Reyes, J. C., Histone availability as a strategy to control gene expression. RNA Biol 2017, 14 (3), 281–286.

